# Virus-mediated transient expression techniques enable genetic modification of *Alopecurus myosuroides*

**DOI:** 10.1101/2020.01.28.923466

**Authors:** Macarena Mellado-Sánchez, Faye McDiarmid, Victor Cardoso, Kostya Kanyuka, Dana R. MacGregor

## Abstract

Even though considerable progress has been made in weed ecology, weed molecular biology has been hindered by an inability to genetically manipulate weeds. Genetic manipulation is essential to demonstrate a causative relationship between genotype and phenotype. Herein we demonstrate that virus-mediated transient expression techniques developed for other monocots can be used in black-grass (*Alopecurus myosuroides*) for loss- and gain-of-function studies. We not only use virus induced gene silencing (VIGS) to create the black-grass exhibiting reduced *PHYTOENE DESATURASE* expression and virus-mediated overexpression (VOX) to drive GREEN FLUORESCENT PROTEIN, we demonstrate these techniques are applicable to testing hypotheses related to herbicide resistance in black-grass. We use VIGS to demonstrate that *AmGSTF1* is necessary for the resistant biotype Peldon to survive fenoxaprop application and show the heterologous expression of the *bialaphos resistance* gene with VOX is sufficient to confer resistance to an otherwise lethal dose of glufosinate. Black-grass is the most problematic weed for winter-cereal farmers in the UK and Western Europe as it has rapidly evolved adaptions that allow it to effectively avoid current integrated weed management practices. Black-grass also reduces yields and therefore directly threatens food security and productivity. Novel disruptive technologies which mitigate resistance evolution and enable better control over this pernicious weed are therefore required. These virus-mediated protocols offer a step change in our ability to alter genes of interest under controlled laboratory conditions and therefore to gain a molecular-level understanding of how black-grass can survive in the agri-environment.

**One Sentence Summary:** Virus-mediated transient expression techniques create loss- and gain-of-function mutations in black-grass and show causation between specific genotypes and measurable changes in herbicide resistance.

## Introduction

Black-grass is an agricultural weed that requires novel innovative control strategies. This weed is a real threat to crop productivity and food security as it competes with crops and reduces yields (Naylor, 1972, Naylor, 2003, Moss et al., 2016, Cook and Roche, 2018). Black-grass has found ways to overcome both herbicides and cultural practices and therefore circumvents the weed control practices currently available to farmers. In the early 1980’s black-grass had demonstrable resistance to most of the graminicides appropriate for use in cereals (Moss and Cussans, 1985). Since then, multiple-herbicide resistance has become widespread in the UK and Western Europe (Délye et al., 2007, Hicks et al., 2018, Heap, 2020). A single black-grass plant can produce thousands of seeds which are shed before harvest and germinate after the next crop is sown. As the lifecycle of black-grass completes between cultivations, it ensures that the seed bank is replenished annually, and the infestation both persists and increases. Each year since 1990, more hectares of land have been treated for black-grass (Hicks et al., 2018). Increased treatments result in increases costs as when weeds become herbicide resistant, farmers spend more money on control (Service, 2013). There are cultural management options which can effectively reduce black-grass populations (Doyle et al., 1986, Allen-Stevens, 2017). However, not all farmers can or choose to adopt the cultural controls that are required for complete control of black-grass as such integrated pest management practices usually require significant input of time, major changes in infrastructure or farming practice, or loss of income during the process (Oakley and Garforth, 1985, Moss, 2019). Therefore, new methods are urgently required to control this costly weed that reduces yield.

To be able to design and deploy sustainable and effective weed management strategies, is critical to understand how black-grass circumvents our current control practices. To gain this understanding, we need to know what genes are underpinning black-grass’s success as an agricultural weed. If we are to demonstrate that we truly understand the gene(s) that control a given phenotype, we need to do hypothesis-led research where we demonstrate causation between genotype with phenotype. This type of hypothesis-led research requires a means to alter the expression of specific targets and assess their phenotypic consequences – i.e. we need to be able to genetically manipulate black-grass.

Transient transformation techniques offer the means to specifically alter gene expression *in planta* in a low-to medium-throughput manner within timeframes that are relevant for researchers and farmers. Of the transient techniques that are available (reviewed in Jones et al. (2009) and Canto (2016)), virus-mediated transient expression techniques offer many advantages. With these techniques the viral genome is modified to heterologously express or to induce RNA-interference to silence a gene of interest. There are many different virus vectors that have been adapted for use *in planta* (Robertson, 2004, Lee et al., 2015a). Once introduced into the plant via rub inoculation, the virus vector multiplies and spreads to new leaves and new tillers within the plant replicating itself and therefore expressing the foreign sequences of interest it carries (Lindbo et al., 2001). Viral vectors can be used to induce loss-of-function through native RNA-interference or posttranscriptional gene silencing pathways, or to promote expression of the heterologous protein of interest (Lindbo et al., 2001). These are viral induced gene silencing (VIGS) or viral induced over-expression (VOX) respectively.

Herein, we demonstrate that the viral vector systems such as those based on *Barley Stripe Mosaic Virus* (*BSMV*) and *Foxtail Mosaic Virus* (*FoMV*) that work well in wheat and several other cereal crops (Lee et al. 2012, Lee et al., 2015b, Bouton et al., 2018) can be adapted to induce gain-or loss-of-function of specific genes in black-grass. These genomic technologies are ideal for functionally validating genes of interest under laboratory conditions and with these we have a unique opportunity to directly alter gene expression in black-grass and thereby functionally validate black-grass genes, including those that underpin its weedy traits.

## Results Summary

Here we demonstrate that both VIGS driven by *BSMV* (Lee et al., 2015b) and VOX driven by *FoMV* (Bouton et al., 2018) can be used successfully in black-grass (Figures 1-3). We demonstrate efficient silencing of *PHYTOENE DESATURASE* (*PDS*) and heterologous expression of GREEN FLUORESCENT PROTEIN (GFP, Figure 1). These two visible markers are often used to demonstrate the VIGS and VOX are functioning in a given species (e.g. Hiroaki et al., 2012, Lee et al., 2012, Lee et al., 2015b, Bouton et al., 2018, Gunupuru et al., 2019). The molecular data we have gathered indicates that the virus-mediated techniques result in the appropriate loss-or gain-of-function at the cellular level (Figure 1). We also demonstrate that VIGS and VOX are suitable to testing hypotheses related to herbicide resistance exhibited in two different populations (Figures 2-3). Our data supports previous conclusions drawn from chemical inhibition studies (Cummins et al., 2013) but moreover, directly demonstrate in black-grass that *AmGSTF1* is necessary for the archetype resistant biotype Peldon to resist 1.5x field rate fenoxaprop (Figure 2). We can also use VOX to provide resistance to an otherwise lethal dose of glufosinate by heterologously expressing the *bar* resistance gene (Figure 3).

**Figure 1:**
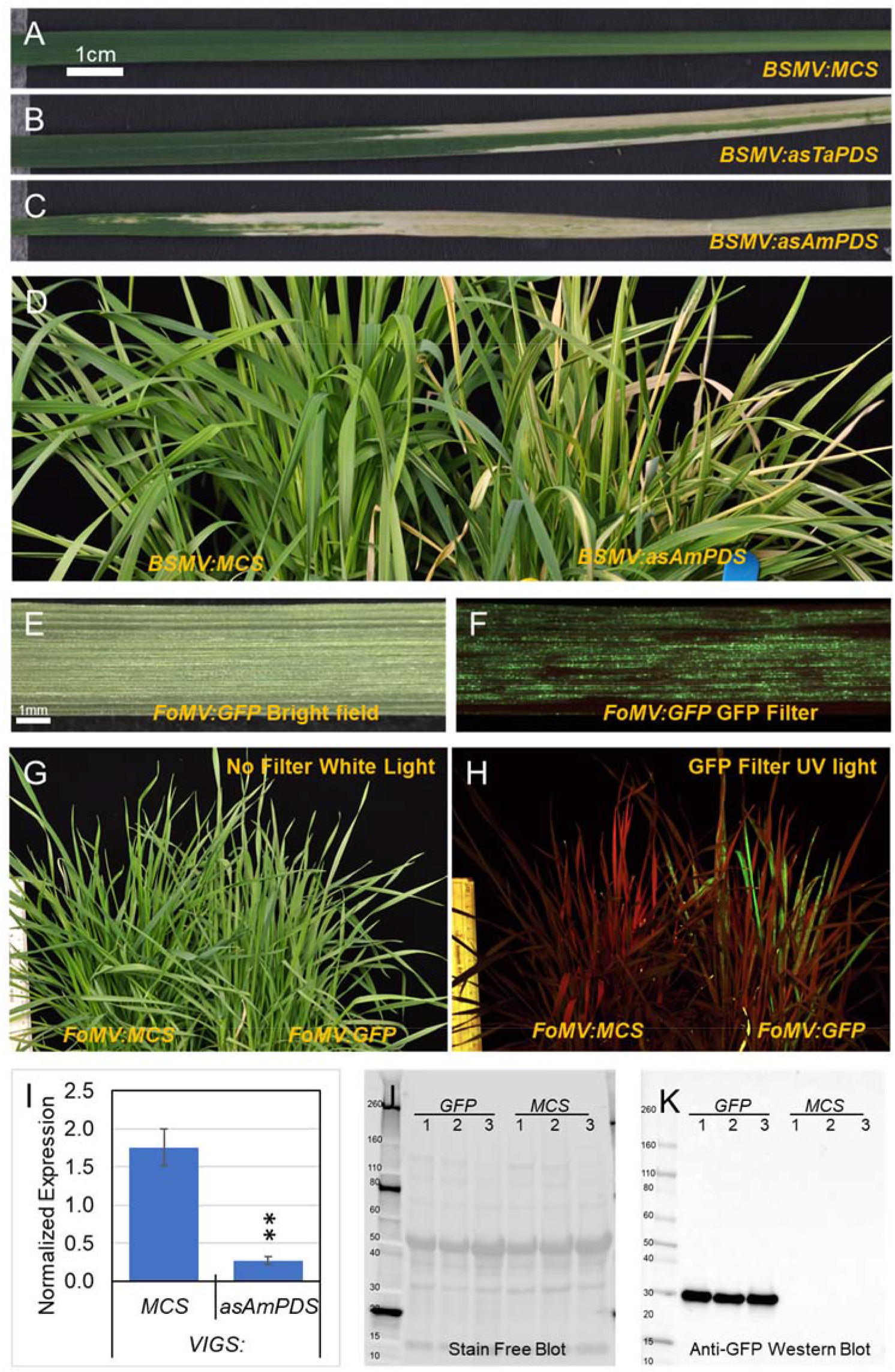
Virus induced gene silencing (VIGS) and virus mediated overexpression (VOX) are possible in black-grass. Data are representative of at least three independent replicates. A-C) Phenotypes of black-grass (Peldon) leaves that have been infected with Barley Stripe Mosaic Virus (*BSMV*) carrying either A) an empty multiple cloning site (MCS), or the MCS with a portion of *PHYTOENE DESATURASE* (*PDS*) in antisense from either B) wheat (as*TaPDS* from Lee et al., 2015) or C) black-grass (as*AmPDS*). D) Whole plant phenotypes of plants from A or B infected with *BSMV:MCS* or *BSMV*:*asAmPDS* as labelled. (E-F) Phenotypes of black-grass (Peldon) leaves that have been infected with Foxtail Mosaic Virus (*FoMV*) carrying *GREEN FLUORESCENT PROTEIN* (*GFP*) from Bouton et al. (2018) under either E) bright field microscopy or F) using the GFP3 filter set. G-H) Phenotype of whole black-grass (Peldon) plants that have been infected with *FoMV:GFP* photographed using a Nikon D90 illuminated with E) white light and no filter or F) blue light using a Dual Fluorescent Protein flashlight through a long pass filter. G-H) Whole plant phenotypes of plants in E & F infected with *FoMV:MCS* or *FoMV:GFP* as labelled and photographed through G) white light and no filter or H) blue light using a Dual Fluorescent Protein flashlight through a long pass filter. I) qRT-PCR of *PDS* normalised against *UBIQUITIN* (*UBQ*) in Peldon plants inoculated with *BSMV:MCS* or *BSMV:asAmPDS*. The data are averages and standard errors from five independent biological replicates each. Asterix indicates a significant difference between that treatments using a Student’s T-Test with * indicating P > 0.05 and ** P > 0.01 compared to the *BSMV:MCS* treated samples. J) Stain free blot showing total protein extracted from Peldon plants inoculated with *FoMV*:*GFP* or *FoMV:MCS* as labelled. Three independent protein extractions per treatment are shown. The size of the bands on the ladder are indicated. K) The blot shown in J probed with anti-GFP followed by Anti-Rabbit IgG–Peroxidase antibody and ECL analysed on a CHEMIDOC MP Imaging Instrument using the manufacturers specifications for optimal and automated acquisition.

**Figure 2:**
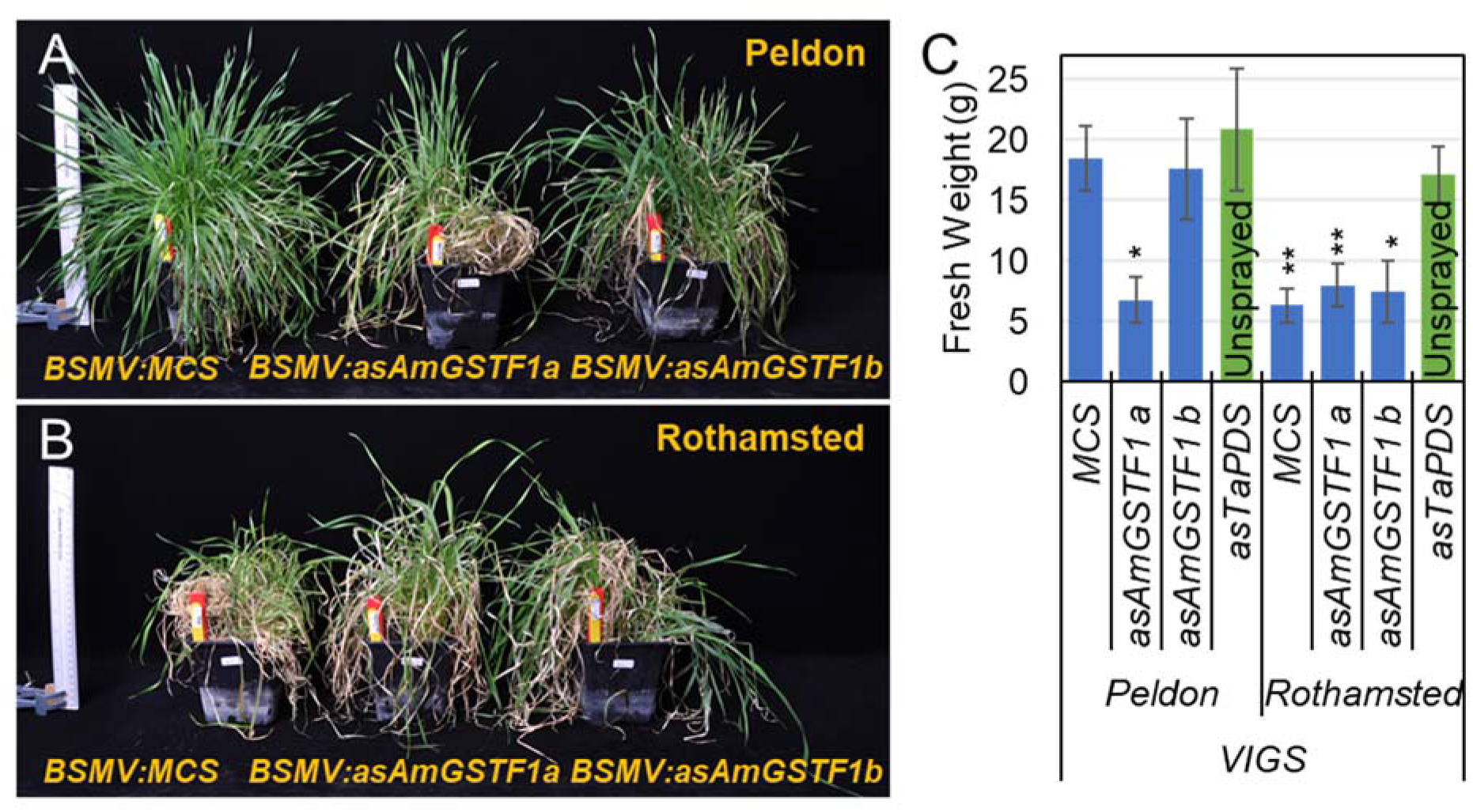
Altering *AmGSTF1* expression using Barley Stripe Mosaic Virus (*BSMV*) is sufficient to revert Peldon herbicide resistance to levels comparable to Rothamsted. Data are representative of three independent replicates. A & B) Phenotypes of Peldon (A) and Rothamsted (B) plants infected with *BSMV* with an empty multiple cloning site (MCS), or two different 200 bp regions of *AmGSTF1* in the antisense direction (*AmGSTF1a* from 6 to 205 bp after the start codon, *AmGSTF1b* from 321 to 520 bp after the start). Photographs were taken 3 weeks after treatment with 1.5x field rate fenoxaprop. C) Fresh weights of greater than 10 plants per treatment in figures A and B taken at 4 weeks after treatment with 1.5x field rate fenoxaprop. Averages and standard errors are shown. Asterix indicates a significant difference between that treatment and the unsprayed plants using a Student’s T-Test and * indicating P > 0.05 and ** P > 0.01.

**Figure 3:**
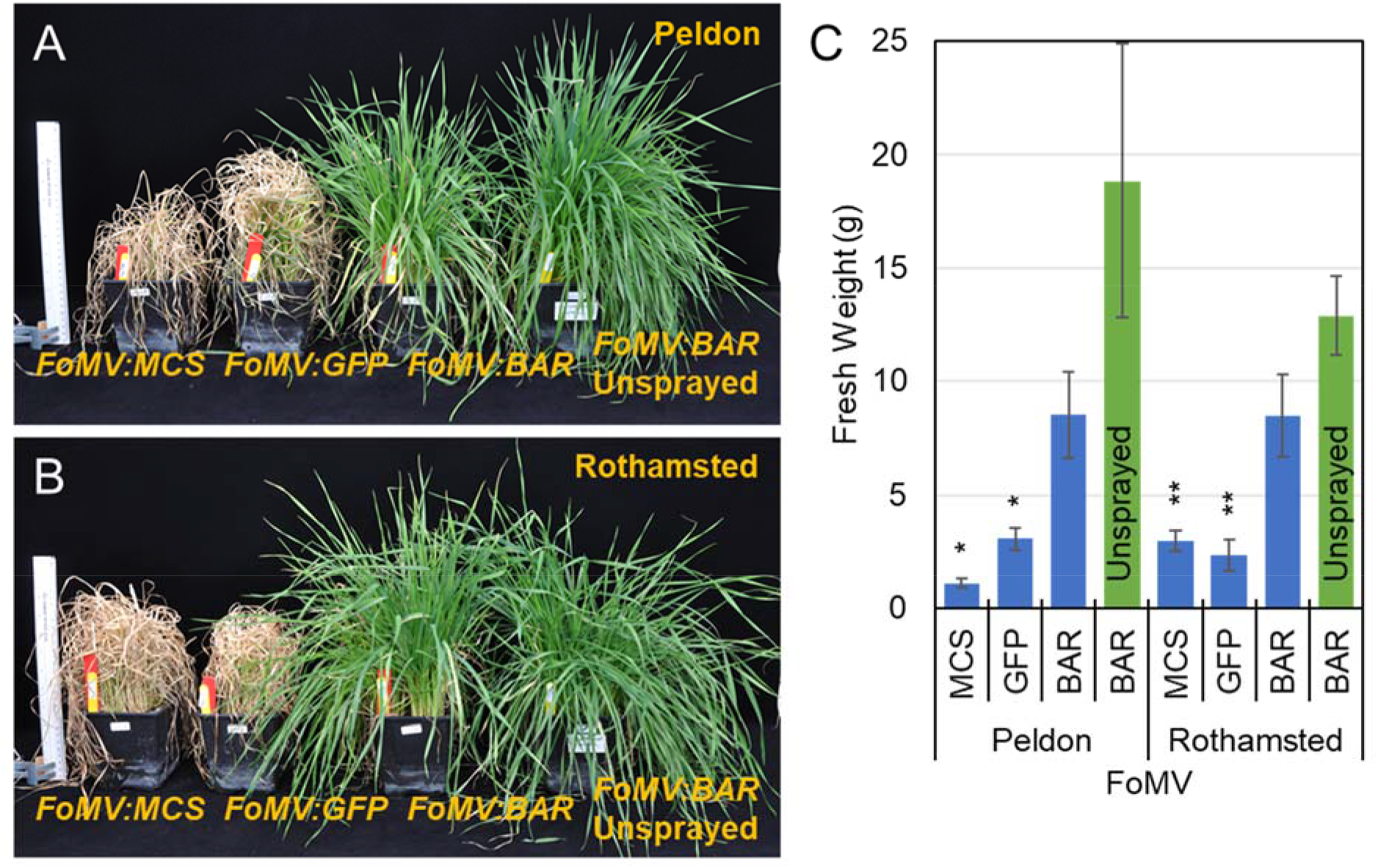
Inoculation with Foxtail Mosaic Virus (*FoMV*) carrying the *bar* resistance gene is sufficient to confer resistance to 0.5% Challenge 60® in Peldon or Rothamsted plants. Data are representative of three independent replicates. A & B) Phenotypes of Peldon (A) and Rothamsted (B) plants infected with *FoMV* carrying an empty multiple cloning site (*MCS*), or the MCS with *GREEN FLUORESCENT PROTEIN* (*GFP*) or bar (bar). Unless indicated by “unsprayed”, plants were treated with 0.5% Challenge. Photographs were taken 2 weeks after treatment. C) Fresh weights of 9 or more plants per treatment (only 5 plants in the case of *FoMV:bar* unsprayed) in Figures A and B taken at 2 weeks after treatment with 0.5% Challenge. Averages and standard errors are shown. Asterix indicates a significant difference between that treatment and the unsprayed plants using a Student’s T-Test and * indicating P > 0.05 and ** P > 0.01.

## Results

In order to determine whether virus induced gene silencing (VIGS) was possible in black-grass, we targeted for silencing *PHYTOENE DESATURASE* (*PDS*) using the wheat *Barley Stripe Mosaic Virus* (*BSMV*) vector system (Lee et al., 2012, Lee et al., 2015a, Lee et al., 2015b). Black-grass plants infected with *BSMV* carrying an empty multiple cloning site (*BSMV:MCS*; Figure 1A) show no outward signs of viral infection. When plants were infected with *BSMV:asTaPDS* (Lee et al., 2012, 2015b), they showed clear loss of green colour within 5-11 days post inoculation (Figure 1B). Similar results were obtained using a vector containing an equivalent portion of the *PDS* gene isolated from black-grass cDNA (Figure 1C). This loss of colour corresponds with a decrease in *AmPDS* RNA as measured by qPCR (Figure 1I). Interestingly, the loss of *AmPDS* phenotype was stable and persisted in the tillered plants (Supplemental Figure 1). Therefore, infection with the established vector carrying a portion of the wheat (Figure 1B) or black-grass (Figure 1C) *PDS* gene in antisense orientation is sufficient to induce loss of green colour as predicted.

In order to determine whether virus mediated overexpression (VOX) was possible in black-grass, we used the *Foxtail Mosaic Virus* (*FoMV*) system developed by Bouton et al. (2018) to attempt heterologous expression of GREEN FLUORESCENT PROTEIN (GFP) in black-grass. When black-grass was inoculated with the *FoMV:GFP* there were no visible symptoms of virus infection in black-grass leaves under bright field imaging (Figure 1E). However, using GFP filters and excitation lamp, GFP fluorescence was clearly visible in the same leaf (Figure 1F). At the level of the whole plant, there were no obvious differences between plants treated with *FoMV:MCS* and *FoMV:GFP* when viewed under white light (Figure 1G), however GFP-specific fluorescence was obvious in the *FoMV:GFP* treated plants when a Dual Fluorescent Protein flashlight and a long pass filter were used (Figure 1H). Fluorescence was visible in treated plants from 9-14 days post inoculation onwards. Autofluorescence (here red) from plants treated with *FoMV:MCS* or *FoMV:GFP* can also be seen (Figure 1H) as not every leaf is manifesting the phenotype. The presence of the GFP protein was confirmed by Western blot analysis (Figures 1J & K). While total protein content and banding patterns were similar between samples taken from plants infected with *FoMV:GFP* and *FoMV:MCS* treated plants (Figure 1J), a band of the appropriate size was detected only in the protein preparations from *FoMV:GFP* treated plants when anti-GFP antibody was used (Figure 1K). Unlike *BSMV* treatments, there was no evidence that the phenotypes generated by *FoMV* treatment can be propagated through tillering (Supplemental Figure 2). When tillers that were visibly exhibiting GFP fluorescence (*n*=4 of Peldon or *n*=13 Rothamsted) were transplanted, none of them showed fluorescence 12, 15, 19 or 23 days later (Supplemental Figure 2).

Using *BSMV* or *FoMV*, the virus vector-induced phenotypes did not manifest in every cell or every leaf of inoculated plants (Figure 1). For example, some of the cells (Figure 1D) and leaves (Figure 1F) of plants infected with *FoMV:GFP* appeared red (due to chlorophyll fluorescence) when viewed under UV or a blue light with the appropriate filters. Gain-or loss-of-function is dependent on the presence of the virus (Ruiz et al., 1998) and the viruses move from cell to cell from the inoculated tissues into developing tissues (Petty et al., 1990, Lawrence and Jackson, 2001), therefore tissues existing before the inoculation will not be infected and will not exhibit the desired phenotypes. These observations are typical of virus-mediated techniques (Singh et al., 2018). However, it is important to note as the herbicides used herein (Figures 2 & 3) were contact foliar sprays.

The data in Figure 1 are presented from the archetype herbicide resistant biotype “Peldon”. The archetype herbicide sensitive biotype “Rothamsted” displayed equivalent phenotypes when infected with *BSMV:asAmPDS* or *FoMV:GFP* (Supplemental Figure 3 & 4).

As stated above, there are currently no methods for weeds that allow for the relationship between genotype and phenotype to be tested. Frequently in black-grass literature, the glutathione transferase AmGSTF1 has been implicated in non-target site herbicide resistance (NTSR) in black-grass. It has demonstrated ability to detoxify herbicides *in vitro* and its protein concentration is correlated with the level of herbicide resistance manifested by plants in the wild, while in the laboratory, chemical inhibition of it reverses herbicide tolerance and heterologously expressing the *AmGSTF1* gene coding sequence in *Arabidopsis* is sufficient to confer herbicide resistance (Cummins et al., 1997, Cummins et al., 1999, Brazier et al., 2002, Dixon et al., 2002a, Dixon et al., 2002b, Reade and Cobb, 2002, Skipsey et al., 2005, Menchari et al., 2007, Cummins et al., 2011, Cummins et al., 2013, Tétard-Jones et al., 2018). For the first time, VIGS allows for us to determine if there is direct causation between AmGSTF1 and herbicide resistance in the black-grass biotypes of interest. The biotype Peldon has long been known for its ability to survive herbicide treatments (Moss, 1990). The plants treated herein were from a purified population, meaning they were the progeny of individuals that exhibited NTSR herbicide resistance but did not carry any known target site resistance mutations bulk crossed in containment greenhouses. In comparison, Rothamsted is the sensitive biotype that have never been exposed to herbicides (Moss, 1990), and the plants used here were similarly purified in glasshouses from clones of individuals demonstrated to exhibit high sensitivity to all tested herbicides. As expected the purified biotype Peldon plants survived higher doses of fenoxaprop than purified biotype Rothamsted plants (Supplemental Figure 5). Where significance is mentioned, supporting P values from Student’s T-tests are given in Supplementary Table 1

In order to determine whether VIGS of *AmGSTF1* would alter the ability of black-grass plants to resist 1.5x field rate of fenoxaprop, we constructed two *BSMV* vectors with each carrying different 200-bp portions of the black-grass *AmGSTF1* coding sequence in antisense. These regions were chosen using siRNA Finder software (http://labtools.ipk-gatersleben.de/) which identifies sequence regions predicted to produce high numbers of silencing-effective siRNAs. Infecting plants with *BSMV:asAmGSTF1a* (but not *BSMV:asAmGSTF1b*) was sufficient to decrease Peldon’s ability to survive herbicide treatment (Figure 2). The fresh weights of Peldon plants treated with *BSMV:asAmGSTF1a* were similar to herbicide-treated Rothamsted plants (Figure 2). Plants of both biotypes inoculated with *BSMV:MCS* exhibited the expected phenotypes at three (Figures 2A & B) and four weeks (Figure 2C) after treatment with 1.5x field rate of fenoxaprop – Peldon survived whereas Rothamsted died. All Rothamsted plants were significantly affected by the 1.5x field rate fenoxaprop treatment regardless of *BSMV* vector with which they were inoculated (Figures 2B & C). Within a biotype, the *BSMV:asAmGSTF1b* did not alter the phenotype compared to treatment with *BSMV:MCS* (Figure 2). However, treatment of Peldon plants with *BSMV:asAmGSTF1a* resulted in a dramatic increase in the number of dead leaves per plant compared to treatment with *BSMV:MCS* (Figure 2A). There was also a significant corresponding loss of fresh weight after 1.5x field rate of fenoxaprop (Figure 2C). In fact, the fresh weight of the Peldon *BSMV:asAmGSTF1a* treated plants was not statistically significantly different from Rothamsted plants treated with *BSMV:MCS* (Figure 2).

We demonstrate here that it is possible to use VOX to give resistance to an otherwise lethal dose of glufosinate by heterologously expressing the *bialaphos resistance* (*bar*) gene from the *FoMV* vector (Figure 3). We cloned the *bar* gene into the *FoMV* vector as this vector holds larger inserts than *BSMV* and was successfully used in other monocots for overexpression and functional analysis of visual markers and apoplastic pathogen effector proteins (Bouton et al., 2018). *bar* encodes for a PHOSPHINOTHRICIN ACETYLTRANSFERASE and was originally isolated from *Streptomyces hygroscopicus* (De Block et al., 1987). The bar enzyme acetylates the active isomer of the glufosinate-ammonium herbicide, and its systemic expression using virus vector thereby provides tolerance to foliar application of this otherwise non-selective herbicide; *bar* has been used to safely provide herbicide resistance to transgenic plants since the late 1980s (Hérouet et al., 2005). Both Peldon and Rothamsted biotypes are susceptible to glufosinate treatment, albeit with different ED_50_ (Supplemental Figure 5). For all the applications, a dose that was lethal to both biotypes was applied. Plants inoculated with *FoMV:MCS*, or those inoculated with *FoMV:GFP* and exhibiting visible fluorescence, all died within two weeks after application of glufosinate (Figure 3). This is apparent due to high numbers of dead leaves on the plant (Figures 3A & B) and a significant reduction in fresh weight (Figure 3C). When either Peldon or Rothamsted plants were inoculated with *FoMV:bar*, although they were clearly affected by the glufosinate treatment, they were dramatically greener (Figures 3A & B) and had fresh weights that were not statistically different from unsprayed *FoMV:bar* plants (Figure 3C) two weeks after application of herbicide. Similar to the evidence above for *FoMV:GFP*, we have no evidence that resistance to glufosinate conferred by *FoMV:bar* is able to persist through tillering (Supplemental Figure 2). With these data we demonstrate that VOX with the *FoMV* vector is suitable for gain-of-function analyses in black-grass relating to herbicide resistance.

## Discussion

Although progress has been made in understanding weed ecology, advancement in understanding weed molecular biology has been impeded by the lack of molecular genetics tools. Most notably, the lack of methods to genetically modify weeds, has meant that no functional validation of genes of interest directly in this plant species was possible till now. The results presented herein demonstrate that cause and effect studies correlating genotype with phenotype are now possible in black-grass. Our results demonstrate that both loss-(Figures 1 & 2) and gain-of-function (Figures 1 & 3) analyses are possible, allowing for questions of necessity and/or sufficiency to be addressed. Not only were we able to recapitulate the standard controls for loss-and gain-of-function analysis using VIGS and VOX with the appropriate molecular support (Figure 1), we have also demonstrated that these techniques allow us to effectively address hypotheses regarding whether a specific gene is necessary or sufficient to confer herbicide resistance in black-grass. These functional genomics techniques worked well for the black-grass populations widely recognized by the field as the archetype resistant (Peldon) and sensitive (Rothamsted) biotypes. Therefore, we can do experiments that compare within and between biotypes. Our data demonstrate that *AmGSTF1* is necessary for Peldon’s enhanced ability to survive fenoxaprop treatment (Figure 2) and that *bar* is sufficient to confer glufosinate resistance to both Peldon and Rothamsted biotypes (Figure 3).

As far as we are aware, this is the first report of functional gene analyses in this agriculturally important weed species and as such, the VIGS and VOX techniques established here offer a step change in the type of questions that can now be asked in weed biology. *FoMV* can hold much larger target sequences than *BSMV* (Bouton et al., 2018), and as both vectors are capable of inducing loss-of-function or gain-of-function (Lee et al., 2012, Lee et al., 2015b, Liu et al., 2016, Bouton et al., 2018) this offers flexibility regarding the length of the coding sequences that can be used and the hypotheses that can be tested. Having a transient treatment is advantageous as it unlinks the field season from the research; this is particularly important for the study of weeds where approximately two thirds of the most problematic weeds globally are single-season or annual weeds with reproductive cycles that are tightly linked with the outside environment (Zimdahl, 2018).

Gene silencing through VIGS offers many useful possibilities. Many of the genes thought to underpin non-target site resistance in several different weeds act through increased expression in the resistant biotype of enzymes that detoxify xenobiotics (Cummins et al., 1999, Cummins et al., 2013, Gaines et al., 2014, Laforest et al., 2017, Yang et al., 2018). Therefore, to reverse herbicide resistances of this sort, the ability to induce loss-of-function of specific genes is required. The fact that both *BSMV* and *FoMV* can deliver posttranscriptional virus-induced gene silencing opens new technical possibilities as the foreign inserts accepted by these viruses and the behaviour of them differs (Lee et al., 2015b, Liu et al., 2016). For instance, the *BSMV*-induced phenotypic change persists through tillering (Supplemental Figure 1) while the *FoMV*-induced changes did not (Supplementary Figure 2). The ability to tiller transgenic plants opens the potential for clonal analyses to be done. Tillered plants are required for creating dose response curves, testing different herbicides, or quantifying life history traits (e.g. (Comont et al., 2019)). Since black-grass is an obligate allogamous species (Sieber and Murray, 1979) with high genetic diversity and low genetic differentiation (Menchari et al., 2007), tillering appears to offer the only opportunity to compare like genotypes directly to like.

Our data indicate that the *AmGSTF1a* in antisense in the *BSMV* vector (Figure 2) was able to alter herbicide resistance in Peldon through post-transcriptional gene silencing. Therefore, we directly demonstrate that *AmGSTF1* is necessary for Peldon’s ability to survive fenoxaprop. These data support previous demonstrations that pre-treatment of Peldon plans with the suicide inhibitor 4-chloro-7-nitrobenzoxadiazole (NBD-Cl; (Ricci et al., 2005)) enhances the phytotoxicity of the herbicide chlorotoluron through inhibition of AmGSTF1 (Cummins et al., 2013). The important caveat to the previous data is that NBD-Cl is known to target other GST family members as well as AmGSTF1 (Ricci et al., 2005, Luisi et al., 2016) and therefore is not a clear demonstration that AmGSTF1 alone is required. Although the possibility exists that our results are a consequence of off-target VIGS, the 200bp portion that was effective (Figure 2) was predicted to have low homology to other sequences. In VIGS, the virus delivers the double stranded RNA, which is recognized and cleaved by the Dicer RNase III enzyme generating the 21–23 nucleotides small interfering RNAs that are loaded into the Argonaute endonuclease (Lu et al., 2003, Baulcombe, 2004). Argonaute with its siRNA is guided to cleave the complementary viral RNA or the homologous endogenous plant RNA sequence(s) based on these 21-23 nucleotide sequences (Lu et al., 2003, Baulcombe, 2004). Although this system is ideal for silencing a single specific gene, non-specific gene silencing or off-target silencing can occur when sufficient sequence homology allows the siRNA generated for intended target gene to the degrade mRNA of genes that are not the intended silencing targets (Senthil-Kumar and Mysore, 2011a). This could be advantageous as it is therefore possible to do multiple gene VIGS repression for redundant or conserved genes. This has been shown to be possible in *Nicotiana benthamiana* against two genes involved in starch degradation (George et al., 2012). Chemical inhibitors that broadly disrupt glutathione synthase activity (Cummins et al., 2013) or cytochromes P450 (Elmore et al., 2015) can also alter herbicide tolerance, and therefore the activity of more than one specific gene may be required for resistance. Taking advantage of the ability to express different lengths of the coding sequence in *FoMV* or *BSMV* will allow us to explore these different possibilities.

Likewise, heterologous gene expression driven by *BSMV* or *FoMV* is equally useful as it allows for questions of sufficiency to be addressed. VOX will create single-gene, dominant mutations. Amplification of the *5-enolpyruvlyshikimate-3-phosphate synthase* (*EPSPS*) gene has been reported to confer resistance to glyphosate in *Amaranthus palmeri* populations (Gaines et al., 2010) and these increased copies are hosted on an extrachromosomal circular DNA molecule carrying the EPSPS gene (Koo et al., 2018) without a corresponding change in ploidy (Culpepper et al., 2006). Therefore, VOX techniques would allow us to recapitulate this mechanism of resistance and directly demonstrate that increased expression or copy number is sufficient to confer resistance in the weed species of interest.

These cause-and-effect analyses are not limited to the study of herbicide resistance; as long as the phenotype can be accurately measured, and the virus effect is induced at the right developmental stage, we can determine if altering the expression of any gene of interest results in the expected phenotypic change. There is also the potential that the silencing effects of *BSMV* treatment could be transmitted to subsequent black-grass progeny through seed as there is precedent for this in other species (Bruun-Rasmussen et al., 2007, Jackson et al., 2009, Senthil-Kumar and Mysore, 2011b, Bennypaul et al., 2012). As far as we are aware, there are no reports of *FoMV*-conferred phenotypes being able to be passed to subsequent progeny. Implementing cultural control practices, such as planting spring crops, can reduce black-grass populations (Doyle et al., 1986, Allen-Stevens, 2017, Varah et al., 2019). However, as black-grass has a demonstrated ability to rapidly adapt to chemical control strategies, it is also probable that it has the capacity to overcome cultural controls by rapidly adapting life history traits that allow it to continue to mimic the crop. VIGS and/or VOX targeted against these key life history traits will be useful to understand the potential for contemporary evolution in response to anthropogenic selection by agricultural weed management.

## Conclusion

In summary, VIGS and VOX offer myriad and unparalleled possibilities for doing hypothesis-led research and functionally validating genes of interest in black-grass. Of main importance will be to apply these techniques to do single gene analysis regarding how black-grass is able to circumvent chemical controls, and thereby to gain a molecular level understanding of what allows it to be such a successful weed. Herein we show that AmGSTF1, a protein previously identified through different approaches as playing a role in non-target site resistance (Cummins et al., 1997, Cummins et al., 1999, Brazier et al., 2002, Dixon et al., 2002a, Dixon et al., 2002b, Reade and Cobb, 2002, Skipsey et al., 2005, Menchari et al., 2007, Cummins et al., 2011, Cummins et al., 2013, Tétard-Jones et al., 2018) is necessary for Peldon’s ability to survive 1.5x field rate of fenoxaprop (Figure 2). We also show that it possible to give black-grass resistance to other herbicides when the resistance genes are known; this was done by heterologously expressing the *bar* gene, which was sufficient to confer resistance to glufosinate (Figure 3). Therefore, VIGS and VOX provide a unique opportunity to do hypothesis led research demonstrating causation between specific genotypes and measurable phenotypes in black-grass.

## Materials and Methods

### Plants and growth conditions

*Nicotiana benthamiana* plants were grown in Levington® Advance F2+S Seed & Modular Compost + Sand Compost, in a controlled environment room with 16 h photoperiod, 23 – 20 °C (day – night), 130 μmol m^−2^ s^−1^ light intensity and 60% relative humidity.

For VIGS and VOX Black-grass (*Alopecurus myosuroides)* plants were grown in a controlled environment room with a 16h photoperiod, 26.7 – 21.1 °C (day – night) temperature, 220 μmol m^−2^ s^−1^ light intensity and 50% relative humidity. Seeds from purified blackgrass biotypes Rothamsted “herbicide sensitive” and Peldon “herbicide resistant” were used. These were pre-germinated on two filter-papers in Petri dishes. The filter papers were wetted with 2g/L potassium nitrate. 5-7 days later, 2 or 5 germinated seeds that had a similar sized radicle were chosen to be transplanted to square 11cm pots filled with Rothamsted Standard Compost Mix (75% Medium grade (L&P) peat, 12% Screened sterilised loam, 3% Medium grade vermiculite, 10% Grit (5mm screened, lime free), 3.5kg per m^3^ Osmocote® Exact Standard 3-4M, 0.5kg per m^3^ PG mix, ~ 3kg lime pH 5.5-6.0 and 200ml per m^3^ Vitax Ultrawet). To aid establishment, propagator lids covered seedlings for 2 days following transplantation.

For glufosinate treatment, 0.5 grams of seed were sown within the top 5 cm of Weed Mix (80% Sterilised Screened Loam, 20% Grit (3-6mm Screened, Lime Free), and 2.0kg Osmocote Exact 5-6 month per m^3^) into containers and allowed to grow to three-leaf stage before application of glufosinate.

The seed lines used were from “purified populations”; this is defined as the population that is create when plants specifically selected for the phenotype are allowed to bulk cross in isolation. For Peldon these individuals exhibited strong NTSR herbicide resistance but did not carry any known TSR mutations or for Rothamsted they were the clones from plants confirmed to be sensitive to all herbicides tested.

### Images of leaves and/or plants

Individual leaves were scanned using a Cannon LiDE110 flatbed scanner. Whole plants were photographed with a Nikon NRK-D90(B) camera (serial number 7051046) with elinca sa CH-1020 D-Lite 2 softbox lamps (serial number e/M2 003658 Renes Switzerland) and Velour Vinyl black backdrop (Superior Seamless 234312).

For microscopy, a Leica M205 FA stereomicroscope with Leica DFC 310FX digital camera using LAS AF software (Leica Microsystems, Milton Keynes, UK) was used with white light and no filter or UV illumination and a GFP3 filter set (excitation filter: 470 ± 40nm; emission filter: 525±50 nm) as indicated. Pictures were taken and quantified using Leica LAS AF software (Leica Microsystems Ltd).

For photographs and monitoring of whole plants for GFP fluorescence, a Dual Fluorescent Protein flashlight (Nightsea, Lexington, MA, USA) was used for illumination and visualisation was done through a long pass (510 nm) filter (Midwest Optical Systems, Palatine, IL, USA).

### Extraction and Quantification of RNA

To obtain the cDNA, the entire plant was frozen and ground in liquid nitrogen from which 100 mg used for total RNA extraction using E.Z.N.A.® Plant RNA Kit (Omega Bio-tek). The RNA was converted into cDNA using SuperScript IV RT (Invitrogen cat# 18090010) and Oligo(dT)_20_ Primer (Invitrogen cat# 18418020) with RNaseOUT™ Recombinant Ribonuclease Inhibitor (Invitrogen cat# 10777-019).

Carried over DNA was removed by off-column treatment with RQ1 RNase-free DNase (Promega cat# M610140). qPCR was done using an Applied Biosystems 7500 Fast Instrument with Quantitation -Standard Curve experimental type and Takyon Low ROX SYBR 2X MasterMix blue dTTP (Eurogentec cat# UF-LSMT-B0710) using three-step protocol for optimal sensitivity and 45 cycles in total. Data were normalised to two different control genes: *UBQ* validated by Petit et al., (2012) and against the *UBQ10* (AT4G05320) homologue with primers designed for this study. As similar results were seen, only one control gene is shown. Primers used herein are listed in Supplementary Table 2.

### Extraction and Quantification of GFP Protein

Protein extraction was done using protocols detailed in Gould et al. (2013) and quantification using protocols in Bouton et al. (2018) using BioRad’s ChemiDoc V3 Western Workflow. Total protein was extracted from individual Peldon leaves from plants treated with *FoMV:MCS* or *FoMV:GFP* using HEPES extraction buffer 1 (100 mM HEPES, 20 mM MgCl2, 1 mM EDTA, 0.10% Triton, 20% Glycerol, 2 mM DTT, with final pH of 8.0 with HCl and 10 µl per ml of Protein Inhibitor Cocktail) which was added to the frozen tissue along with two titanium balls. Samples were ground using a Retsch bead mill and centrifuged for 5 min at 4°C. The supernatant was removed to a new tube and centrifuged again. This clarified supernatant was diluted 1:2 and 1:4 in HEPES extraction buffer.

Equal volumes of protein were loaded onto pre-cast BioRad Mini-Protean TGX precast Gel 4-20% (15-wells BioRad cat# 4568096). The gels were run according to the manufacturer’s guidelines (110Volts for 3 minutes, then 250Volts for 20 minutes). Whole protein detection was carried out using stain free imaging settings for Coomassie stain equivalent in the ChemiDoc Imaging System (cat# 12003153). Protein transfer was accomplished using BioRad Trans-Blot® Turbo™ Mini Nitrocellulose Transfer Packs (cat# 1704158) and the Trans-Blot Turbo Transfer System (cat# 1704150) using the manufacturer’s guidelines (2.0A 25Volts for 7 minutes). Once transferred, total protein was imaged using the stain free settings for Ponceau equivalent in the ChemiDoc Imaging System (cat# 12003153). The membrane was also stained with Ponceau S solution (0.1% (w/v) Ponceau S in 5% (v/v) acetic acid) as a loading control and blocked for one hour in 5% Marvel TBST at room temperature. All washes were done three times for 5 minutes with TBST. The primary antibody Invitrogen anti-green fluorescent protein rabbit IgG fraction (cat # A11122) was diluted 1:2000 in 5% Marvel TBST and left to incubate overnight at 4°C. The secondary antibody, anti-rabbit IgG (Sigma cat# A0545) was diluted 1:10,000 in 5% Marvel TBST and left to incubate for 1.5 hours at room temperature. Use of these antibodies to detect *FoMV:GFP* had previous been described in Bouton et al. (2018). Clarity Western ECL substrate (BioRad cat# 1705060) was then applied for visualisation. Detection of chemiluminescence was accomplished using ChemiDoc Imaging System (cat# 12003153) and the on-board image-acquisition software with auto-exposure settings appropriate to the Clarity ECL.

### Cloning of *BSMV* and *FoMV* vectors

The *BSMV:MCS* and *BSMV:asTaPDS* vectors and the methods required to create the black-grass *BSMV:asAmPDS* and *BSMV:asAmGSTF1* variants are described in detail in Lee et al. (2015b) with the following changes. Phusion ® High-Fidelity DNA polymerase (NEB cat# M0530) was used on cDNA libraries made from Peldon plants described above (Extraction and Quantification of RNA) with the primers detailed in Supplementary Table 2. PCR products were gel purified with either Wizard® SV Gel and PCR Clean-Up System (Promega cat# A9281) or Isolate II PCR and Gel Kit (Bioline cat# BIO-52059) according to the manufacturer’s protocols. The target sequences were then cloned into the *BSMV* RNAγ vector *pCa-γbLIC* (*BSMV:MCS*) vector from Yuan et al. (2011) via the ligation independent cloning protocols exactly as described in Lee et al. (2015b). They were fully sequenced using primers in the viral backbone to verify the products. For *BSMV:asAmPDS*, both the Peldon and Rothamsted sequences were cloned. Both were shown to induce the white leaf phenotype; the results shown here are all from the Peldon sequence.

The *FoMV:MCS* and *FoMV:GFP* vectors are published (Bouton et al., 2018). The primers used for the creation of *FoMV:bar* are in Supplementary Table 2. These introduce a NotI site at the 5’ end, mutated TGA stop codons to TAA, and introduced an XbaI site at the 3’ end of the sequence. The *bar* gene was amplified with Phusion ® High-Fidelity DNA polymerase (NEB cat# M0530) from the pAL156 vector (Amoah et al., 2001) and was subcloned into Zero Blunt ® TOPO PCR cloning kit for sequencing (Invitrogen cat# 450159). Once sequencing confirmed there were no PCR-induced mutations, the *bar* gene was removed with the NotI and XbaI restriction sites created via the PCR primers and inserted into the *FoMV:MCS* via traditional cut and paste cloning with T4 DNA ligase (Fisher cat# EL0014) following the manufacturer’s protocols and the ligated products were transformed into JM109 Competent Cells (Promega cat# L2005).

Once the *BSMV* and *FoMV* vectors were confirmed by sequencing, they were transformed into *Agrobacterium tumefaciens* strain GV3101 through standard electroporation techniques and recombinants selected based on survival of dual selection with kanamycin and gentamycin. Individual colonies were selected, multiplied, and verified by colony PCR with the appropriate primers (Supplementary Table 2).

### Preparation of the virus inoculum from *Nicotiana benthamiana*

The recombinant *BSMV* and *FoMV* viruses were propagated via agroinfiltration in *Nicotiana benthamiana* using protocols detailed in Lee et al. (2015b). The leaf that was infiltrated was harvested 3-5 days after infiltration for *BSMV* vectors and 5-7 days after infiltration for *FoMV* vectors. One leaf from three different *N. benthamiana* plants were weighed into foil packets and plunged into liquid nitrogen before being stored at −80°C.

### Rub-inoculation of black-grass

These protocols are based on those published in Lee et al. (2015b) and Bouton et al. (2018) with minor changes. Black-grass seedlings were grown at 27°C day / 21°C night with 16 hours of daylight for 18-29 days or until the 2 or 3 tiller stages. The second leaf on a thick tiller of each plant was chosen for rub-inoculation. To facilitate inoculation, each leaf was marked with a paint-pen, then Carborundum (Technical, SLR, Extra Fine Powder, ~ 36µm (300 Grit), Fisher Chemical cat 10345170) was applied through a cheesecloth to evenly coat the adaxial side of the leaf. The inoculum was prepared by grinding the three agroinfiltrated *N. benthamiana* leaves in a 2:1 (w/v) ratio in 10mM potassium phosphate buffer pH 7. The thumb and forefinger of a gloved hand were dipped in the inoculum and rubbed the length of the leaf 10 times firmly. The plants were incubated in the controlled environment room overnight (covered to create low light conditions) and were returned to standard growth conditions the following day.

### Herbicide Applications and Assessments

Herbicides were applied 14 days after rub-inoculation. The herbicides applied, Fenoxaprop (Foxtrot) or Glufosinate (Challenge), are both commercially available. Treatments were fenoxaprop (Foxtrot, 69 g/l (6.9 % w/w) fenoxaprop-p-ethyl, Headland Agrochemicals) and glufosinate (Challenge-60™, 200 g/L glufosinate-ammonium, Bayer). Fenoxaprop was applied at 1.5x field rate (103.5 g/l of fenoxaprop-p-ethyl) diluted in distilled water. For Figure 4 glufosinate was applied with 0.1% Tween in distilled water at 0.5% (0.3g/l of glufosinate-ammonium). For Supplementary Figure 5, 8 doses of Challenge-60™ and 1 Untreated were used (0.0%, 0.05%, 0.1%, 0.2%, 0.3%, 0.4%, 0.5%, 0.75% and 1.0%) in 0.1% Tween in distilled water and applied using the same methods. The herbicide was diluted to the chosen concentration then transferred into a Cooper Pegler CP 1.5 Mini Pro Sprayer bottle which was used to saturate black-grass plants. After application of fenoxaprop and glufosinate, at 28 days and 14 days later respectively, all plants had observations and photographs taken and were harvested for fresh weights of above ground tissue.

## List of Author Contributions

DRM conceived the original idea and formulated the research plan. DRM designed the experiments with input from MM-S and FM. MM-S, FM and DRM performed the experiments. The black-grass specific vectors were cloned by VC and/or DRM into the cloCantoning sites within viral vectors created by KK from Lee et al. (2015b) or Bouton et al. (2018). KK also created the *FoMV:PV101-GFP* vector and provided guidance and support regarding VIGS and VOX protocols and methods. DRM wrote the article with contributions from all the authors. DRM agrees to serve as the author responsible for contact and ensures communication.

## Accessions numbers

Novel DNA sequences identified for AmPDS (MN936109) and AmGSTF1 (MN936108 associated with AJ010454.1) in this paper have been deposited in GenBank (http://www.ncbi.nlm.nih.gov) with the accession numbers listed.

## Acknowledgments

The authors would like to acknowledge the following people. We thank Ari Sadanandom (Durham University) for his support with the N8 Pump Prime Award that funded the preliminary data investigating *BSMV* application in black-grass during DRM’s Fellowship at Durham University. We thank Richard Hull (Rothamsted Research), Laura Crook (Rothamsted Research), and Paul Neve (AHDB) for providing seed of the Rothamsted and Peldon purified lines. David Comont and Claudia Lowe (Rothamsted Research) are recognized for the data regarding Rothamsted and Peldon fenoxaprop sensitivity to fenoxaprop and Marie Lamarre (Agrocampus Ouest) for Rothamsted and Peldon fenoxaprop sensitivity to glufosinate. We thank Sergio Cerezo-Medina (Rothamsted Research) for providing the *Agrobacteria tumefaciens* strain GV3101 and Caroline Sparks (Rothamsted Research) for the plasmid pAL156. Our gratitude goes to Helen-L Martin (Rothamsted Research) for carrying out the herbicide applications and to Graham Shephard (Rothamsted Research) for assistance with photography. We are grateful to Richard Hull (Rothamsted Research), Laura Crook (Rothamsted Research), David Comont (Rothamsted Research) and Paul Neve (AHDB) for useful discussions regarding the work throughout.

## Supplemental Data

Supplementary Table 1: Student’s T-Test P values to support claims of significance or insignificance used throughout the paper.

Supplementary Table 2: Primers used throughout the paper.

Supplemental Figure 1: The loss of green colour correlated to infection with *BSMV:asTaPDS* or *BSMV:asAmPDS* is stable through tillering.

Supplemental Figure 2: *FoMV* phenotype is not stable through tillering.

Supplemental Figure 3: Peldon and Rothamsted plants two weeks after inoculation with *BSMV:MCS* or *BSMV*:AmPDS exhibit clear VIGS phenotype.

Supplemental Figure 4: Peldon and Rothamsted plants two weeks after inoculation with *FoMV:GFP* exhibit clear GFP fluorescence.

Supplemental Figure 5: Dose-response curves for Rothamsted and Peldon biotypes when challenged with A) glufosinate (Challenge) or B) fenoxaprop.

